# A developmental program that regulates mammalian organ size offsets evolutionary distance

**DOI:** 10.1101/2022.10.19.512107

**Authors:** Yuko Shimamura, Junichi Tanaka, Miwako Kakiuchi, Hemanta Sarmah, Akihiro Miura, Youngmin Hwang, Anri Sawada, Zurab Ninish, Kazuhiko Yamada, James J. Cai, Munemasa Mori

**Affiliations:** Columbia Center for Human Development and Division of Pulmonary, Allergy, Critical Care, Department of Medicine, Columbia University Medical Center, New York, NY, USA; Department of Preventive Medicine, Graduate School of Medicine, the University of Tokyo, Tokyo, Japan; Department of Surgery, Johns Hopkins University, Baltimore, MD, USA; Department of Veterinary Integrative Biosciences, Department of Electrical & Computer Engineering Interdisciplinary Graduate Program in Genetics Center for Statistical Bioinformatics, Texas A&M University, College Station, TX, USA

## Abstract

Pigs are evolutionarily more distant from humans than mice, but their physiological organs are closest to humans. The molecular program leading to a more than 1,000-fold increase in organ size in pigs and humans over that of mice across evolution has not been elucidated. We generated large-scale transcriptional landscapes throughout swine lung development. Our cross-species single-cell molecular atlas let us discover swine progenitor identities, stage-specific markers, and a core organ-size regulation program (COSRP), well-conserved in swine and humans but less so in mice. Across eight mammalian species, human COSRP promoters showed higher homologies to evolutionary-distant large animals, including pigs, than evolutionary-close small animals. Our study provides a molecular foundation during swine lung development that unveils animal size regulation conserved in the COSRP promoter, independent of genome-wide evolution. COSRP is a critical paradigm for studying thousands-fold changes in biological sizes in evolution, development, cancer, zoology, respirology, organoids, and biotechnology, particularly human-compatible organ generation.

**One Sentence Summary:** A cross-species developmental molecular atlas identified the indicator of lung and animal size beyond evolution

## Main Text

Evolution changes the size of animals to adapt to the environment(*1*, *2*). Organ size regulation is a fundamental program of the developmental process, however, the molecular program that governs large organ size across evolution remains a mystery(*3*–*5*). None of the technologies achieved functional human internal organ generation because there has been a considerable knowledge gap in how human-compatible organ sizes develop. The search for organ size regulators in specific organogenesis programs in large animals, which show more than 1,000-fold organ size differences compared to small animals, has the great potential to elucidate the hidden mechanisms of organ size diversification in evolution and advance human-compatible organ size bioengineering in various fields (*3*–*5*). In this regard, we focused on the development of the pig lungs because its molecular program has been a complete mystery for centuries, and early embryonic development in pigs still lacks organ primordia and has not yet led to a dramatic increase in organismal size compared to mice.

The lungs are one of the most diversified organs during evolution and the most complex organs to replicate. The need for life-sustaining gas exchange through the lungs facilitated environmental adaptation by evolving the lung’s developmental program over billions of years, allowing species-specific physiological morphologies, surfactant protein production, and unique circulatory systems(*6*, *7*). Conversely, species such as the plethodontid salamander, which breaths through skins, are lungless by evolutionary design(*8*). Mice and swine have been used as essential model organisms for decades, and both harbor life-sustaining, well-lobulated, branched lungs(*9*–*13*). Multiple studies of comparative genomic evolution show that pigs primitively separated around 97 million years ago from mice and humans(*14*–*17*). Mice and humans were divided about six million years later. Thus, swine genome-wide sequences are evolutionarily apart from humans as compared to mice(*14*–*17*). Contradictorily, pigs harbor physiological organs similar to humans, distinct from small rodents by more than 1,000-fold in size(*17*, *18*). Given the prolific feature and the physiological similarities to humans, pigs are considered an ideal resort for xenogeneic organ transplantation and the host animals for human organ bioengineering by blastocyst complementation for the ultimate cure for various refractory diseases(*19*, *20*). Although many signaling pathways, including Hippo-Yap, JNK, and IGF-1(*3*), are involved in organ size regulation, their signaling alone does not explain the regulatory mechanism for more than 1,000-fold difference in organ size between small animals and animals of comparable size to humans. To address this question, we performed comparative cross-species bioinformatic analyses in pig, mouse, and human lung development.

## Results

### Pig embryonic lungs develop exponentially along with their body size

The mouse’s developing lung stage has been molecularly and morphologically well-characterized, whereas swine lung development is rarely characterized (*10*, *13*). To fill the huge knowledge gap on the molecular program on how and when swine lung organ size change is initiated, we harvested developing lungs from miniature pigs harboring human-compatible organ size(*19*) (**Fig. 1a**). Based on the gross morphology, we identified embryonic gestation day18 and 19 (hereafter, E18 and E19) as the primary lung bud formation stage (**Fig. 1b**). Interestingly, the swine primary lung bud exhibited slightly enlarged appearance on E19 than E18. After secondary buds appear on E20~E23, sequential pseudostratified epithelial branching occurs (**Fig. 1b**). Through the pseudoglandular stage, the lungs expand their size exponentially from E40 to E70, along with body weight changes (**Fig. 1c**).

**Figure 1.**
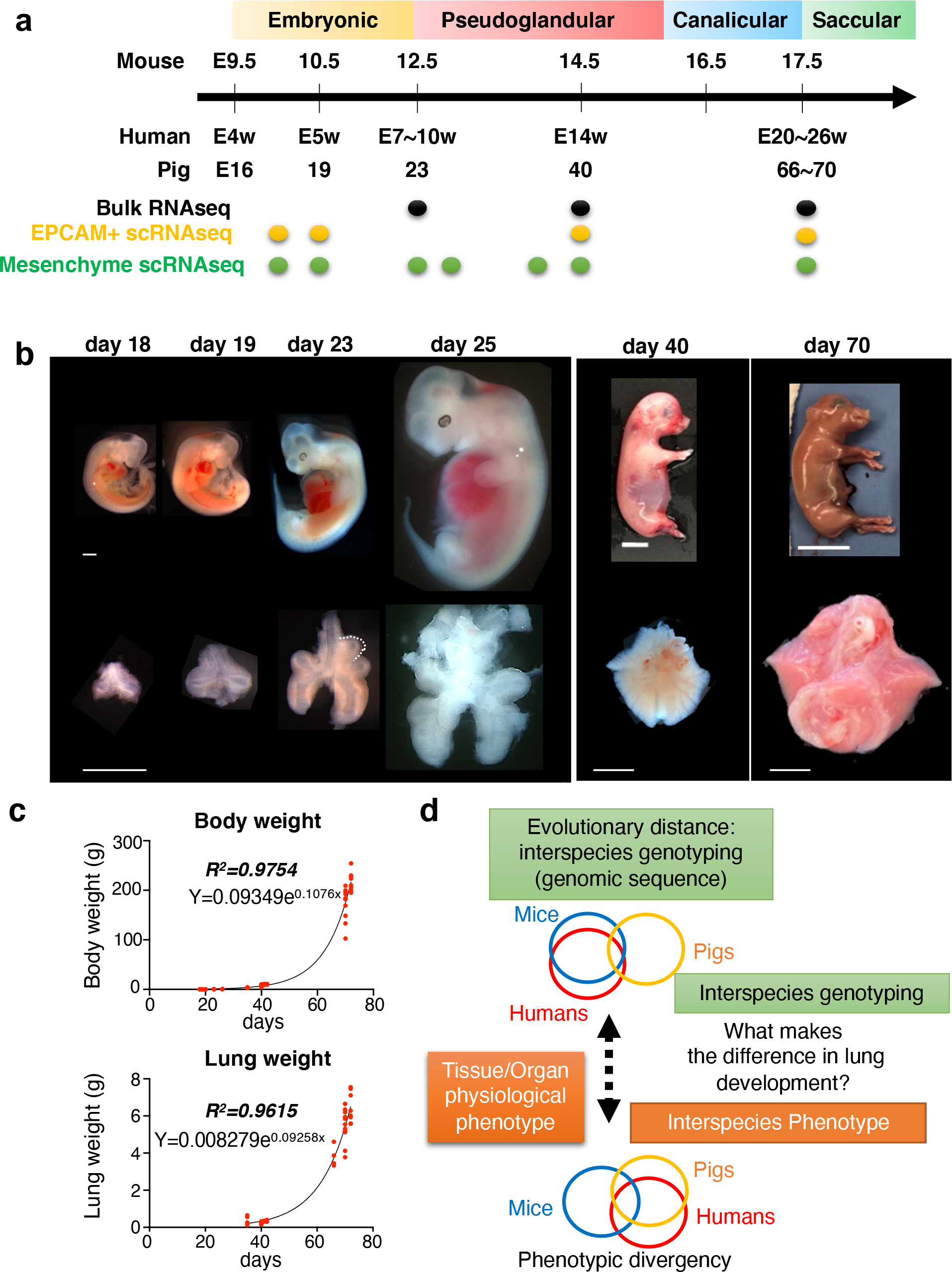
The dramatic change of organismal and lung size during swine development. **(a)** Schematic diagram: Developmental stages and sampling of pig lungs compared to mice and humans. Each experiment is indicated by a dot (Black: bulk RNA-seq, Yellow: EPCAM-sorted epithelial scRNA-seq, Green: mesenchyme scRNA-seq). **(b)** The representative gross morphology of pig embryos at each embryonic gestation period (E) 18, 19, 23, 25, 42, 70. **(c)** Graphs: Exponential change of the weight of pig embryos and lungs. Each red dot indicates each embryonic pig specimen. **(d)** The schematic model depicting the discrepancy between interspecies genotyping and phenotype. The human organ size phenotype is closer to a pig than a mouse, while the human genomic sequence is evolutionarily closer to a mouse than a pig. Scale Bar**s**: 1mm (E18-25), 1cm (E40 embryo and lung, and E70 lung), 5cm (E70 lung)

### Pigs and humans share unique organ size regulator programs, distinct from mice

Although swine are evolutionarily more distant from humans than mice(*14*–*17*), the molecular program required for the acquisition of a human-compatible lung organ size phenotype remains elusive (**Fig. 1d**). We isolated RNA from miniature swine lung tissues at E26, E40, and E70 for bulk RNA-seq analysis (**Fig. 1a**) and defined swine developmental stages by gross lung morphology, histology, and cross-species transcriptome (**Fig. 1a, Extended Data Fig. 1, and Supplementary Table 1**). To extract latent variables underlying phenotypic divergence across species, we performed a principal component (PC) analysis. Using PC1 (39% variance) and PC2 (28% variance), the most extensive variant sets in PC analyses confirmed a relatively uniform sampling without significant batch variance (**Extended Data Fig. 2a**). Importantly, PC3 (13% variance) indicates the developmental time course over the species (**Fig. 2a**). With the combination of PC1 and PC3, the most prominent variance among all PCs, we realized that swine developing lungs have a closer transcriptome variance to humans than mice do (**Fig. 2a**). To identify the genes involved in the human and swine transcriptome variance proximity, apart from mice, we ranked genes based on their contribution to the PC1 minus axis. We performed GO term analysis on the top 10% of genes. Of note, we found the program of anatomical structure size regulation (ASSR) (GO: 0090066) (**Fig. 2b and Extended Data Fig. 2b**). Unexpectedly, the neuronal-like developmental program (GO:0007409, 0099536, 0051960, 0010975, 0051963) highly overlays ASSR. Thus, we defined the neuronal and ASSR programs as the **<u>c</u>**ore of the **<u>o</u>**rgan **<u>s</u>**ize **<u>r</u>**egulation **<u>p</u>**rogram (COSRP) that includes 230 genes (**Fig. 2b**, red circled area). Further, the Venn chart illustrated that the COSRP neuronal-like program overlapped with 57.1% of the ASSR program (80 out of 140 ASSR genes) (**Fig. 2c and Extended Data Fig. 2c**).

**Figure 2.**
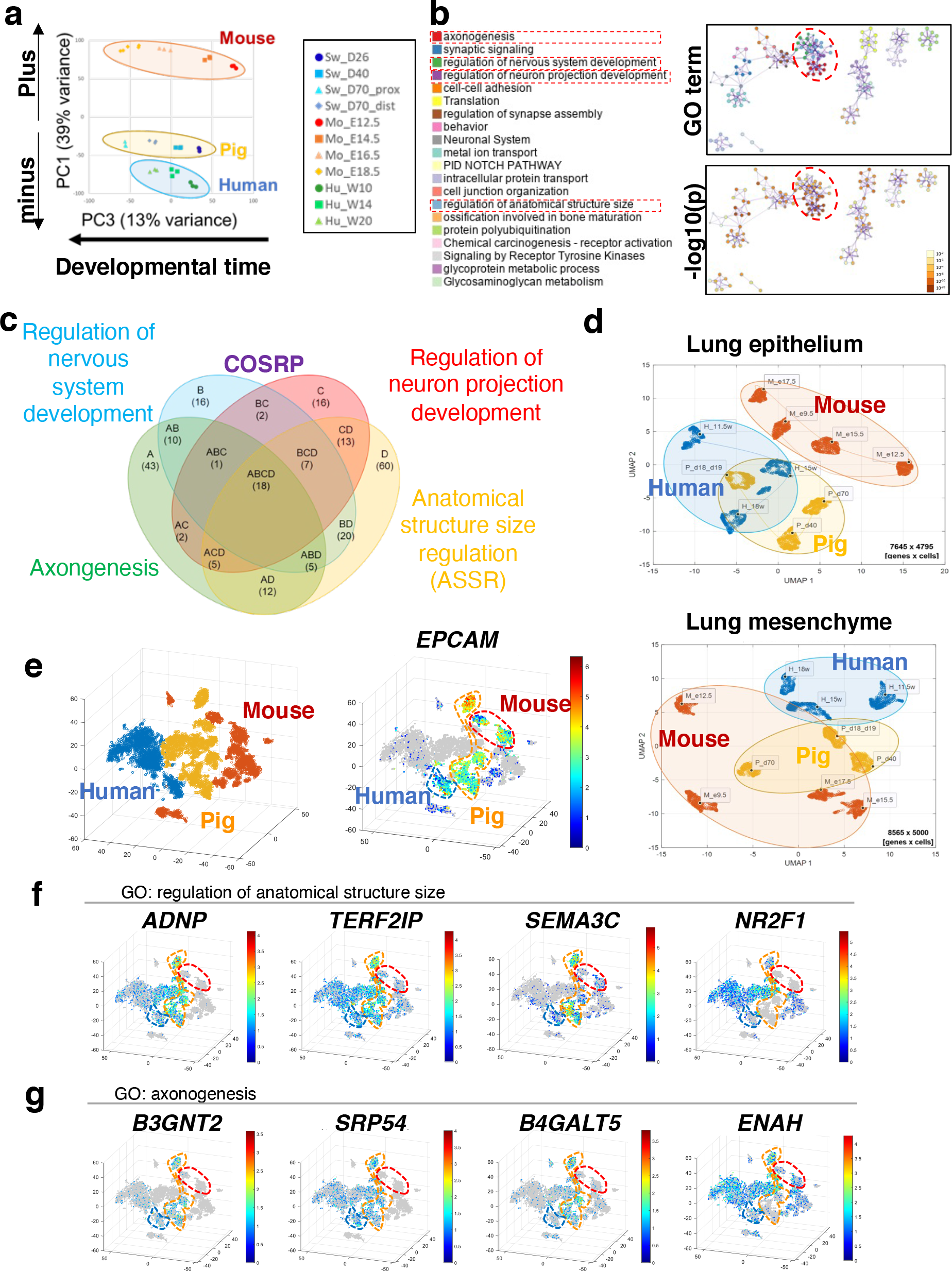
Cross-species transcriptome analyses identify the <u>c</u>ore of the <u>o</u>rgan <u>s</u>ize <u>r</u>egulation <u>p</u>rogram (COSRP) during swine lung development. **(a)** PCA analyses showing human, mouse, and pig embryonic lung transcriptional variance. PC3: the developmental time of each species. PC1 minus in the Y axis: the human and swine had similar variance over time, distinct from mice. **(b)** Gene ontology (GO) terms network analysis visualizing the top 10% genes contributing to PC1 minus axis and GO-term associated network: top: Each color indicates GO term category in the left panel. Bottom: colored by significance (P-value: 10^−2^~10^−20^). Red dotted circle: COSRP. (**c**) Venn chart plot visualizing 45 COSRP genes composed of four GO terms; Axonogenesis, Regulation of nervous system development, Regulation of neuron projection development, and Anatomical structure size regulation (ASSR). **(d)** Cross-species scRNA-seq UMAP: swine transcriptome closer to humans than mice in developing lung epithelium (top) and mesenchyme (bottom) over time. **(e)** 3D tSNE plot: human, mouse, and swine EPCAM^+^ lung epithelium and EPCAM^−^ mesenchyme scRNA-seq combined data. 3D tSNE feature plot for representative COSRP genes: **(f)** *ADNP, TERF2IP, SEMA3C*, and *NR2F1*. **(g)** *B3GNT2, SRP54, B4GALT5*, and *ENAH* were highly enriched in swing and human epithelium or mesenchyme but rarely in mice.

To map which cell type acquires the unique COSRP features conserved in swine and humans but not in mice, we harvested swine-developing lungs (a total of 83,433 single-cell RNA-seq from 11 independent samples; 7 distinct time points for lung mesenchyme and 4 particular time points for EPCAM^+^ FACS-sorted lung epithelium). We utilized SCGEATOOL(*21*) to perform subclustering annotation with combined datasets across time points, as well as data-driven analyses of gene expression patterns, gene-regulatory networks, and trajectory inference. For cross-species comparison, we combined datasets of swine, human, and mouse-developing lungs and analyzed them in an integrated manner (**Fig. 2d**). The UMAP analyses for the transcriptome profile of swine lung epithelium throughout swine lung development revealed relatively closer progenitor identity with that of human lung epithelium than mice (**Fig. 2d**). In the transcriptome profile of the lung mesenchyme, mice were still far apart from humans overall (**Fig. 2d**). Among 230 COSRP genes in the Venn chart (**Extended Data Fig. 2c**), we discovered that 45 genes were expressed in human and swine lung development but rarely in mice by the cross-species 3D-tSNE scRNA-seq analyses (**Fig. 2e**). Unexpectedly, we also discovered that 17 COSRP genes, including the neuronal-like program and ASSR, were ubiquitously conserved in both EPCAM^+^ pig and human lung epithelium or mesenchyme but not well-conserved in mice (**Fig. 2e, f, g, and Supplementary Table 2**). These ubiquitous gene expression patterns during swine and human lung development but rarely in mice showed three distinct distributions: first, 3 COSRP genes were exclusively enriched in the EPCAM^+^ epithelium (*SEMA3C, B3GNT2*, and *B4GALT5*); second, 4 genes exclusively enriched in EPCAM^−^ mesenchyme (*COL5A2*, *ENAH*, *PRKG1*, and *NR2F1*); third, 10 genes were enriched in both, such as (*SRP54, EFNA5, ANTXR1, ADNP, ID4, TERF2LP, ADAMTS7, CCDC85B, TET1*, and *ZFP36L2*) (**Fig. 2e, f, g, Extended Data Fig. 2c, and Supplementary Table 2**). These results suggested that 45 COSRP genes are the responsible organ size regulator conserved in swine and human lung development but rarely in mice over time.

### Unique expression pattern of COSRP genes conserved in swine developing lung epithelium

To uncover which cells express COSRP genes in developing lung epithelium and systematically characterize early lung epithelial progenitor molecular identities, we analyzed scRNA-seq data of swine lung epithelium across four-time points from E18, E19, E40, and E70 (27,287 genes for a total of 9,240 live cells) (**Fig. 3a**). Human lung epithelial progenitors exhibit *SOX2^+^SOX9^+^* double-positive distal lung bud progenitors, never seen in the mice at the pseudoglandular stage(*22*). On E18 and E19 of the primary lung bud formation stage, *SOX2* and *SOX9* were indeed co-expressed almost entirely in the lung epithelium (**Fig. 3b**, asterisks), and confocal immunofluorescent (IF) protein expression analyses further confirmed their distribution and SPB and SPC protein expression in the distal lung bud regions (**Fig. 3c**, asterisks, **and Extended Data Fig. 3a-d**). In contrast, at the beginning of the sacculation stage on E70, SOX2 and SOX9 rarely overlapped (**Fig. 3b, c, d**, arrow). On the other hand, SOX2 and SOX9 coincided with E40 lung buds, but the SOX2 expression level was relatively decreased compared to the SOX2^+^ proximal domain (**Fig. 3b, c, d**, arrowhead).

**Figure 3.**
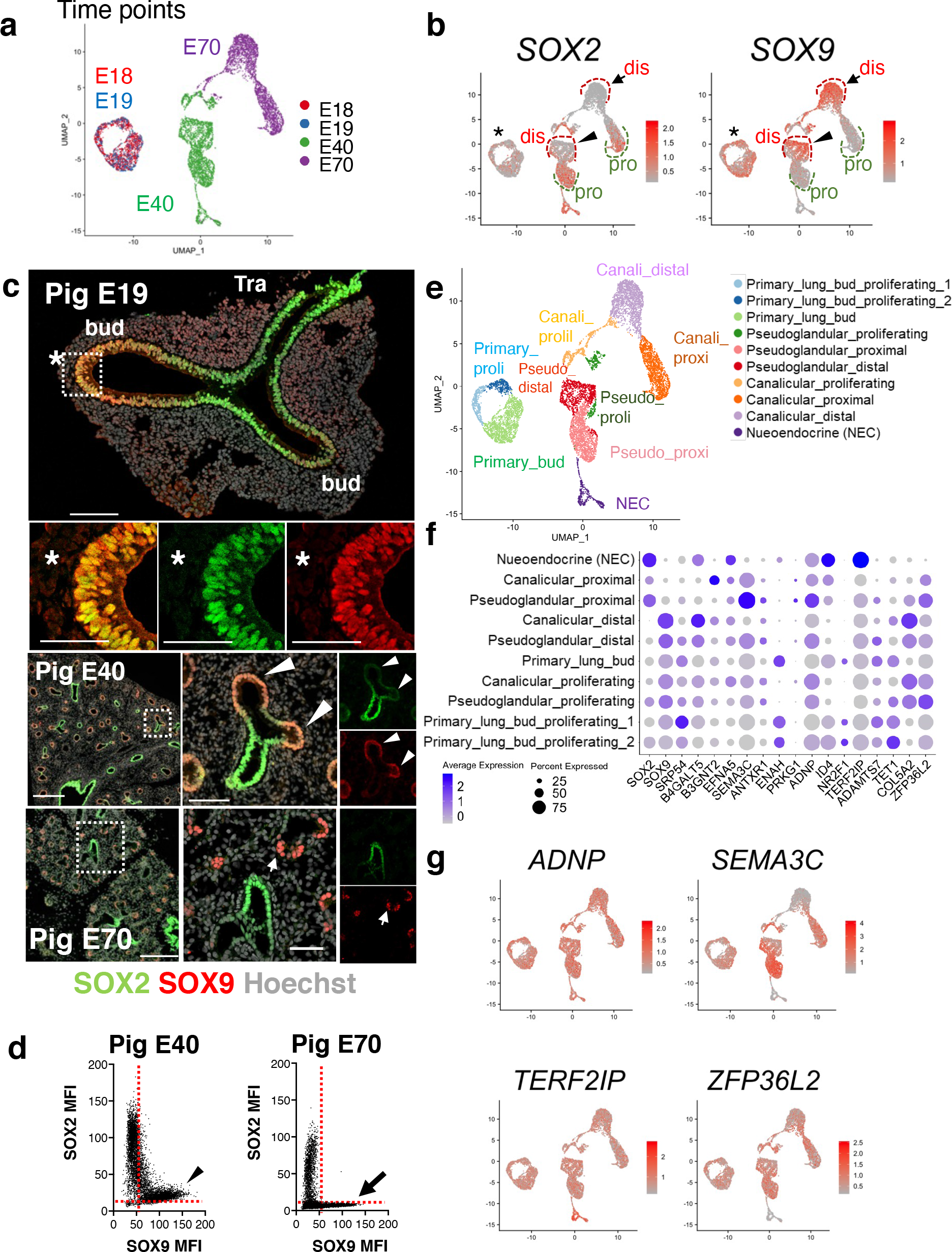
A unique human-like embryonic porcine lung epithelial program. UMAP feature plot for scRNA-seq depicting the samplings from embryonic lung epithelial cells from pig E18, E19, E40, and E70 **(a)**, *SOX2* and *SOX9* gene expression patterns (**b**), ten clusters in swine developing epithelial cells (**e**), and a ubiquitous expression of COSRP genes (ADNP, SEMA3C, *TERF2IP*, and *ZFP36L2)* in swine developing epithelial cells throughout lung development (**g**). **(c)** confocal IF imaging: embryonic pig lungs at E19, E40, and E70: SOX2 (green) and SOX9 (red) were co-expressed on the primary lung buds (asterisks*) and day40 distal lung buds (arrowheads), while only SOX9 were present on the day70 distal lung buds (arrows). **(d)** Graphs: qualitative morphometric analysis for SOX2 and SOX9 protein expression in the nucleus of pig E40 and E70 epithelial cells. Each dot: mean fluorescent intensity (MFI) of Sox2 or Sox9 in a cell nucleus. Arrow; day70 distal epithelial cells, arrowhead: day40 distal epithelial cells. **(f)** Dot plot showing the expression levels of *SOX2, SOX9*, and representative COSRP genes. Dis: distal (red dot lines), Pro: proximal (green dot lines) lung epithelial compartment. Scale bars: top: 100μm, enlarged: 50μm, middle 500μm, enlarged: 100μm, Bottom: 200μm, enlarged: 50μm

ScRNA-seq analysis further identified 10 epithelial clusters (**Fig. 3e**). E18 and E19, the primary lung bud formation stage, showed similar profiles but were utterly different from E40 and E70, pseudoglandular and canalicular lung bud stages, respectively. Using classical mouse lung epithelial markers(*23*), the dot plot and feature plot visualized airway and alveolar epithelial cells during swine lung development (**Extended Data Fig. 3a, b**). Although it is not related directly to COSRP, we also discovered novel stage-specific markers such as *NES, SLC16A3, FABP3*, and *SLC2A3* explicitly in the *SOX2^+^SOX9^+^* double-positive swine primary lung bud epithelial cells while *Slc16a3, Fabp3*, and *Slc2a3* were partially expressed in E9, the mouse primordial lung stage (**Extended Data Fig. 4a, b**). *SLC22A3*, *GPLD1*, *SHISA2*, and *KCNJ15*, were the markers of pseudoglandular~canalicular specific *SOX9^+^* swine lung bud epithelial cells, in which *SLC22A3*, *SHISA2*, and *KCNJ15* were well-preserved in human lung development but not in mice (**Extended Data Fig. 4a, b, c**). Among the 45 COSRP genes, we found 12 genes *(SRP54, B4GALT5, B3GNT2, EFNA5, ANTXR1, ADNP, ID4, TERF2IP, SEMA3, ADAMTS7, CCDC85B, TET1*, and *ZFP36L2*) were moderately or highly expressed in most lung epithelium throughout swine lung development (**Fig. 3f, g, and Supplementary Table 2**). The dot plot analyses showed the spatially distinct expression pattern of COSRP genes in the subclusters (**Fig. 3f**). Briefly, *NR2F1* and *ENAH* were exclusively expressed at the primary lung bud epithelium but rarely later. Interestingly, *B4GALT5, SOCS1, and COL5A2* were highly enriched in the SOX9^+^ distal lung bud. In contrast, *SEMA3C or B3GNT2 was* relatively increased in the proximal epithelial domain on day40 or day70, respectively. *ID4* was enriched in neuroendocrine cells (NEC) (**Fig. 3f, g, and Supplementary Table 2**).

**Figure 4.**
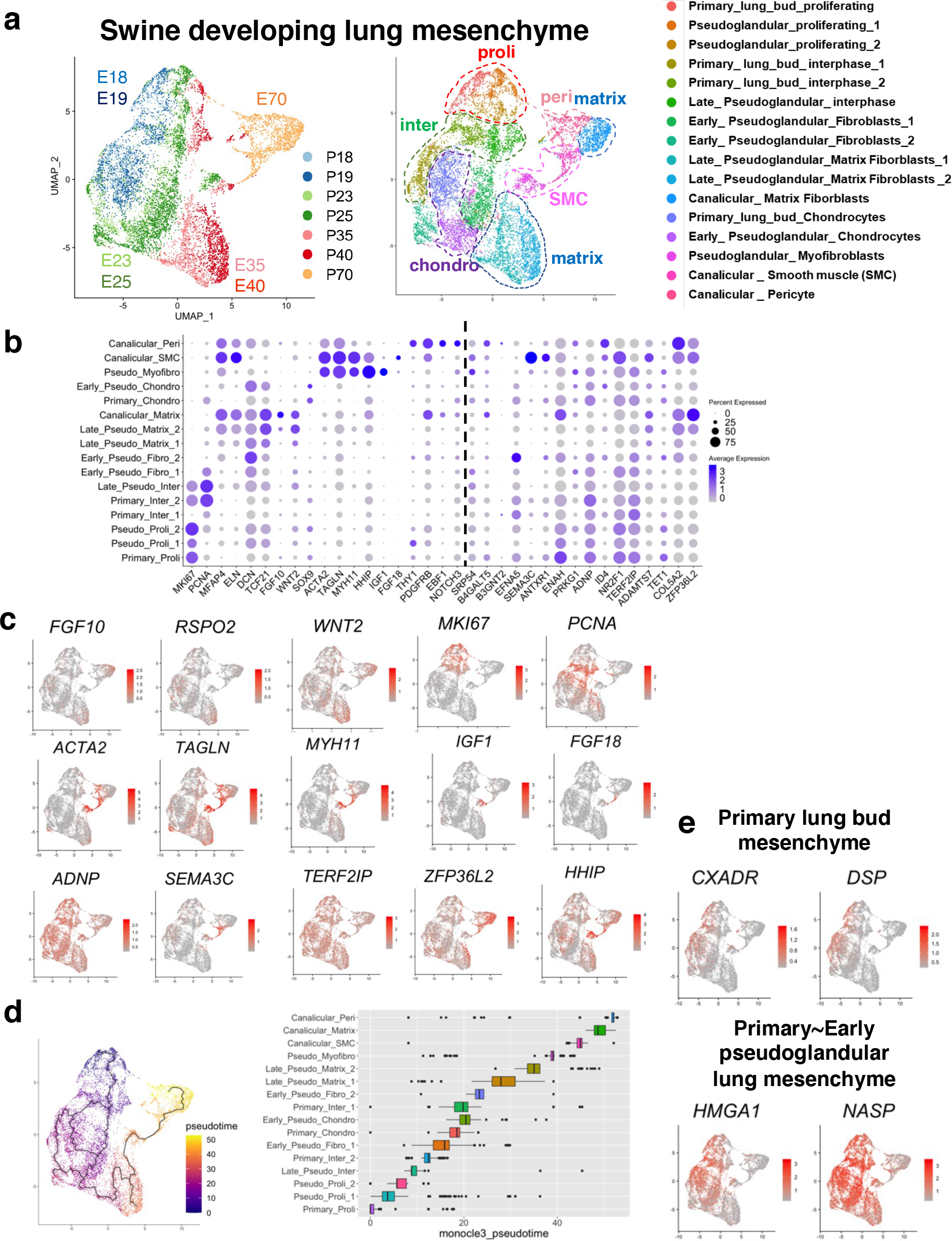
COSRP gene transcriptional profiling of embryonic porcine lung mesenchymal cells. **(a)** UMAP visualization of lung mesenchymal cells from pig E18, E19, E23, E25, E35, E40, and E70. A dot blot (**b**) and UMAP feature plot (**c, e**) for COSRP genes (*ADNP, ENAH, NR2F1, TERF2IP, COL5A2, SEMA3C, and ZFP36L2*) and representative classical lung mesenchymal markers (distal lung bud markers: *FGF10, RSPO2*, lung mesenchyme specifier*: WNT2*, pericyte: *NOTCH3, EBF1, THY1*, smooth muscle cells (SMC) and myofibroblasts*: ACTA2, TAGLN, MYH11, HHIP, FGF18, IGF1*, chondrocyte: *DCN, SOX9, PDGFRB*, Matrix fibroblast*: MFAP4, ELN, TCF21*, proliferating cells (*MKI67, PCNA*))(**c)**, and the stage-specific markers: primary lung bud stage (*CXADR* and *DSP*) and primary to pseudoglandular stage (*HMGA1* and *NASP*) (**e**). **(d)** Monocle pseudotime analysis of embryonic porcine lung mesenchymal cells. Cells are labeled by pseudotime. Boxplot showing the distribution of pseudotime within each cell group.

### Identification of COSRP genes ubiquitously expressed in swine developing lung mesenchyme

To reveal the COSRP gene expression in the lineage trajectory and diversification of developing pig lung mesenchyme, scRNA-seq datasets of whole lungs on E18, 19, 23, and 25 and peripheral lung tissues on E35, E40, and E70 were analyzed using Seurat and classified the cellular subclusters (**Fig. 4a**). We demonstrated that chondrocyte progenitors emerged at the swine primary lung bud stage, marked by *SOX9*. At the E35 and 40 middle pseudoglandular stage, matrix fibroblast and smooth muscle cell (SMC) clusters appeared (**Fig. 4a**). At E70, late pseudoglandular~canalicular stage, SMC, pericyte, and matrix fibroblast clusters were found (**Fig. 4a**). A dot plot and feature plot analysis of swine lung mesenchymal cells showed stage-specific subclusters and their gene expression profiles well-known in mouse studies (**Fig. 4b, c**). For example, *NOTCH3* and *EBF1* were enriched in the vascular pericyte clusters at the canalicular stage. At the swine lung bud stage, a part of proliferative mesenchyme or matrix fibroblast expressed *FGF10*, a critical mitogen for primary lung bud formation and branching morphogenesis(*24*), *WNT2*, a marker for the lung mesenchyme specifier(*25*), *RSPON2*, a feature of lung bud-specific mesenchyme in human lung development(*26*)(**Fig. 4b, c**). *ACTA2, TAGLN, MYH11*, and *HHIP* were highly expressed in the smooth muscle cell (SMC) clusters at the canalicular stage and myofibroblasts at the pseudoglandular stage(*27*). *ACTA2* and *IGF1-positive* myofibroblasts emerged from E23 and differentiated into *ACTA2*- and *FGF18*-positive smooth muscle cells at E70 **(Fig. 4b, c**). Pseudotime analysis revealed stage-specific genes enriched in the primary lung bud fibroblast-specific stage (*CXADR* and *DSP*), primary to pseudoglandular stage (*HMGA1* and *NASP*) (**Fig. 4d, e)**. A matrix fibroblast (MF) labeled by *MFAP4, ELN, MEOX2, TCF21*, and *DCN* has been abundantly enriched in swine and human lung development but rare in mice(*28*). We found *MFAP4*, *ELN*, and *TCF21* gradually increased their expression levels during swine lung development and peaked at E70, most likely by its maturation (**Extended Data Fig. 5a, b**). *TCF21* and *DCN* expressions were conserved across swine and humans but not in mice (**Extended Data Fig. 5b**). We identified novel MF markers through scRNA-seq of swine developing lung mesenchyme. *CAV2*, *RGCC*, and *INMT* were highly expressed in swine MFs. These genes were expressed in human lung mesenchymal tissue but were not abundant in mice (**Extended Data Fig. 5c**). These results suggest that the mesenchymal subcluster of peripheral lung tissue reported in humans is highly conserved in pigs. Among the 45 COSRP genes, 7 *(ERF2IP, COL5A2, ZFP36L2, ENAH, PRKG1, NR2F1*, and *ADNP)* were ubiquitously expressed in the lung mesenchyme throughout swine lung development, but the other COSRP genes showed diverse expression patterns (**Fig. 4c and Supplementary Table 2**).

**Figure 5.**
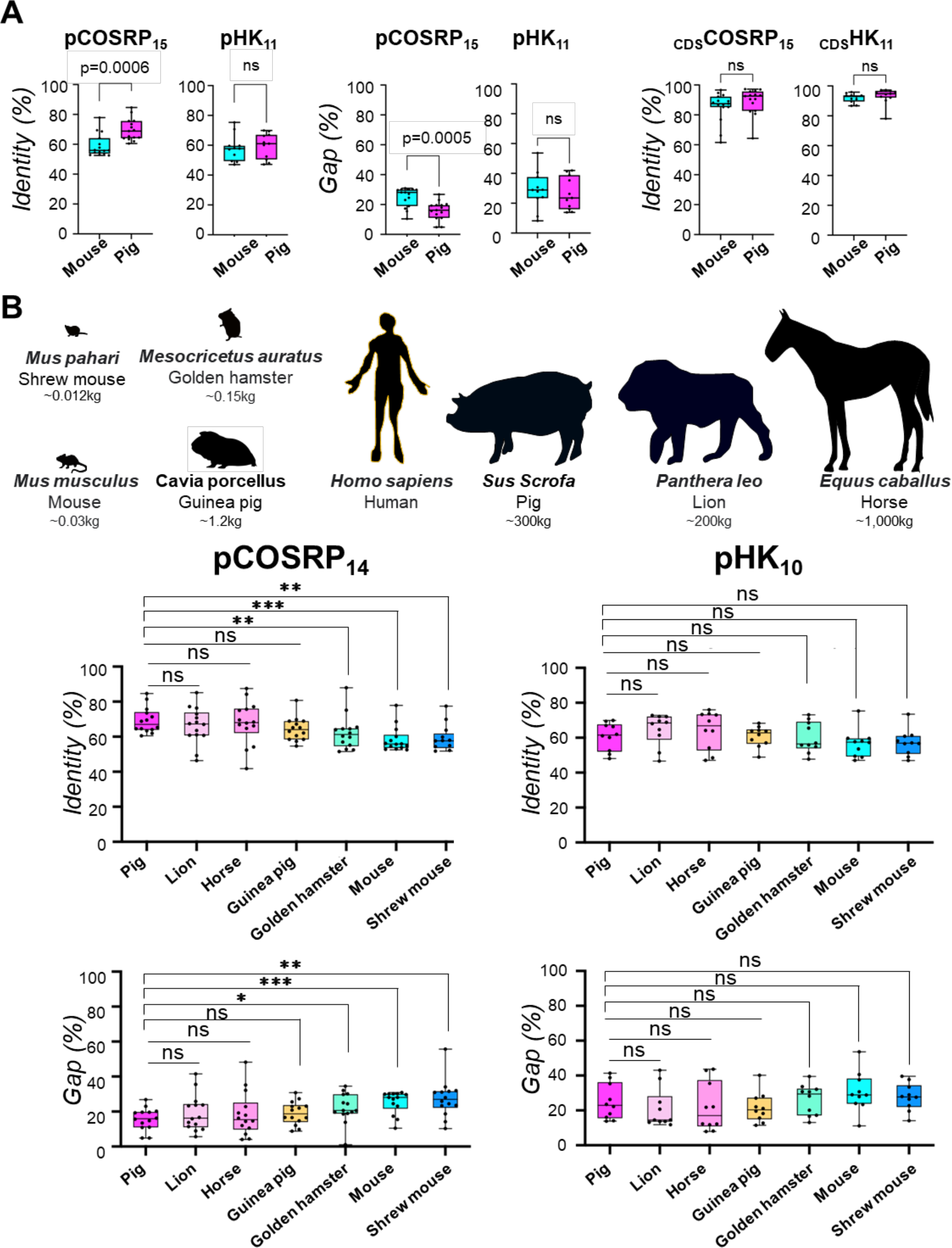
COSRP promoter region encodes a relative organismal size. **(a)** graphs: A box and whisker plot: COSRP or housekeeping (HK) gene promoter homologies (left: % identity, middle: % gap) of mice and pigs to humans. Right: The Coding sequence (CDS) homologies (% identity) of mice and pigs to humans. The homology of the human COSRP promoter region was significantly lower in mice than in swine (p=0.0006, ***P<0.001) while not significant in HK promoter regions or CDS or HK regions. (COSRP gene; n=15, HK genes; n=11) **(b)** top: Schematic diagram of large and small animals with representative body weight. Bottom: Graphs: Homologies (% identity, % gap) of each species COSRP, housekeeping (HK) gene promoters to humans. A dot in each graph (**a, b**) represents each COSRP or HK gene in each species. mean±s.e.m. in 14 COSRP genes or 10 HK genes per group. two-tails Student’s t-test, *P<0.05, **P<0.01, ***P<0.001, ****P<0.0001, ns: statistically non-significant.

### COSRP gene regulatory networks are conserved in swine and humans but not in mice

To elucidate the significant network of cellular functional programs governing the organ size regulators during lung development over mRNA and protein expression patterns, we performed gene regulatory network (GRN) analyses for the 45 COSRP genes. We used swine, human, and mouse lung developmental scRNA-seq databases at each time point by SCGEATOOL(*21*). Based on the scRNA-seq GRN analysis (scGRN), swine lung developmental scGRN showed multiple highly interacting nodes with connected networks similar to humans (**Extended Data Fig. 6a, b**). The highly-dense scGRN network is conserved in swine and human lung development but sparse in mice.

Together, our analyses showed that swine developmental lung’s phenotypic similarity of organ size, cellular phenotype, and scGRN with humans is regulated by COSRP that is not well-conserved in mice. However, it was unclear how organ size regulators represented by COSRP offset the evolutionary distance to humans, in contrast to the mice, which is calculated by each species’ overall genomic sequences(*14*–*17*).

### COSRP genomic promoters are better conserved in large animals than in small animals

Since COSRP mRNA expression depends on its promoter activation, we speculated that swine and humans’ lung organ size phenotypes might rely on the COSRP promoter genomic sequence homologies against the entire genomic evolution. To explore this possibility, we investigated 15 COSRP gene promoter regions (pCOSRP_15_) and 11 HK gene promoter regions (pHK_11_) as controls. To keep the evolutionary significance and avoid the complexity of differential length of promoter analyses across the various species, we analyzed 2kb upstream of TSS: transcription starting site for a human COSRP promoter (pCOSRP) (**Fig. 5a**) (*29*).

Strikingly, we discovered that pCOSRP_15_ aligned sequences showed a significantly higher level of homology between humans and pigs (69.7% ±7.01%) than the homologies between mice and humans (59.1%±7.29%) (P<0.001***). Conversely, pHK_11_ did not significantly (ns: non-significant) differ between the homology comparison with pigs (59.4%±8.16%) and mice (57.9% ±8.35%) (**Fig. 5a**). We also checked the COSRP gene coding sequence (CDS) alignments (_CDS_COSRP_15_). However, _CDS_COSRP_15_ also did not show a significant homology difference in humans between mice (85.3%±9.57%) and pigs (89.1%±8.41%) nor in HK genes (mouse: 91.9%±2.95%, pigs 93.2%±5.23%). Those results indicate that lung organ size regulation depends on the 2kb upstream of TSS for 15 COSRP genes most likely to activate the COSRP program.

COSRP was found by the cross-species bulk RNA-seq analyses when the lung size exponentially increased along with the organismal size’s drastic change (**Fig. 1, 2**). Thus, we hypothesized that pCOSRP homology might be an indicator of lung size but also the organismal size itself among mammalian organisms. Since some organisms were not known the genomic sequence of COSRP or HK promoter homologous regions in the Ensemble, we selected 8 animals (Lion, Pig, Horse, Guinea pig, Golden hamster, Mouse, and Shrew mouse) and used 14 COSRP gene’s promoter regions pCOSRP_14_ and 10 HK genes (pHK_10_) for the homology comparisons (**Fig. 5b**). Strikingly the pCOSRP_14_ genomic homology was well-conserved in mammalian large animals compared to small creatures (**Fig. 5b**). The Pigs (*Sus Scrofa*)’s pCOSRP_14_ homologies (% identity and % Gap) to the humans showed significance to all small animals such as Shrew (*Mus phahari*) (% identity*: p=0.0013*, % Gap: *p=0.0017, respectively*), Mouse (*Mus Musculus*) (*p=0.0006, p=0.005*), and hamster (*Mesocricetus auratus*) (*p=0.0040, p=0.034*), but not to the large animals such as Lion (*Panthera Leo*) (*p>0.05, p>0.05*) or Horse (*Equus Caballus*) (*p>0.05, p>0.05*) (**Fig. 5b**). The pHK_10_ homologies to the humans vary and no significant change between pigs and the other animals (*ns*: non-significant), suggesting pCOSRP_14_ is the unique genomic region highly conserved in large animals but less in small animals.

## Discussion

Our analyses unlocked previously unknown COSRP regulatory mechanisms for lung size along with the body size that contradicts the species’ entire genomic sequence-based evolutionary distance. Since small animals did not show high pCOSRP_14_ homology to humans, even though they are evolutionarily closer to humans than swine, small species most likely needed to lose the pCOSRP_14_ homologies to make their body size smaller to adapt to the environment billions of years ago after the division of pigs and human beings as a survival strategy during evolution. Our results suggest that large organismal size development requires COSRP in developing epithelial and mesenchymal lung progenitors. Further detailed analyses for the COSRP promoters will be required to elucidate which signaling pathways, such as Hippo-Yap, and JNK, coordinate to activate the pCOSRP_14_. Our results indicate that “species” various phenotypic divergence relies more on the responsible promoter element than an entire genomic sequence that determines genome-wide evolutionary distances (*14*–*17*). Thus, our findings provide a novel paradigm for cross-species analysis, organismal size regulations, and lung organ size across the species, distinct from the classical evolutionary phylogenic tree studies. The discovery of COSRP and pCOSRP can be utilized for various studies that will lead to a better understanding of zoology, evolution, pulmonary medicine, regeneration, development, cancer biology, and bioengineering, particularly organ bioengineering compatible with humans.

## Acknowledgments

We thank the members of the Columbia Center for Human Development for the scientific input, members of the Columbia Genome Center for the transcriptional analysis for RNA-seq, the CCTI flow cytometry core (LSRII: NIH S10RR027050), and Columbia Stem Cell Initiative (CSCI) Flow Cytometry core (FACS Area) for FACS and sorting.

## Author contributions

Conceptualization: M.M.

Methodology: M.M., Y.S., J.T., J.J.C., M.K.

Investigation: Y.S., J.T., H.S., A.M., Y.H., A.S., Z.N., K.Y., M.M.

Visualization: Y.S., J.T., M.K., J.J.C., M.K.

Funding acquisition: M.M., J.J.C.

Project administration: M.M.

Supervision: J.J.C., M.M.

Writing – original draft: Y.S., J.T., J.J.C., M.M.

## Data and materials availability

Data available upon request

## Supplementary Materials

Materials and Methods

Extended Data Fig. 1 to 6

Supplementary Table 1 to 2

