## Supplementary information for "A developmental program that regulates mammalian organ size offsets evolutionary distance"

Yuko Shimamura *et al*.

Corresponding authors:
Munemasa Mori,
James J, Cai,

**This PDF file includes:**

Materials and Methods
Extended Data Figure 1 to 6
Captions for Supplementary Table 1 to S2

**Other Supplementary Materials for this manuscript include the following:**

Supplementary Table 1 to S2

### Materials and Methods

#### Animals

Timed-pregnant Yucatan miniature sows were obtained from Sinclair BioResources. We surgically harvested the embryos from E18 to E70 and euthanized the pregnant sows right after the surgery. The body weight of swine embryos and their lungs were measured right after sampling the specimens. The gross morphology images were taken by a digital camera and dissection microscopy (Leica TL5000 elgo, Nikon Labophot 2 microscope). All surgical procedures were performed under the approval of the Columbia University Institutional Animal Care and Use Committee and the USAMRMC Animal Care and Use Review Office (ACURO).

#### Tissue preparation for bulk RNA-Seq

For bulk RNA-seq analyses, we collected the peripheral lungs from E66 and E40 pig embryos, the proximal lungs (around the main bronchus) from E66, and half of the whole lungs from E26. Biological triplicates were prepared from independent embryos (*n=3,* at each time point). Total RNA from those samples was extracted with ReliaPrep RNA Miniprep Systems (Z6111, Promega) according to the manufacturer's protocol. Libraries for RNA-Seq were prepared using Illumina TruSeq chemistry that were sequenced with paired-end sequencing using Illumina NovaSeq 6000. The sequencing reads were performed pseudo alignment to a Kallisto index created from Pig transcriptomes Sscrofa11.1 using Kallisto (0.44.0).

#### Tissue dissociation for scRNA-Seq

Lung buds from E41 and E70 embryos are dissociated with the accutase and stained with DAPI to detect dead cells by FACS analyzer. Live cells were sorted out using FACSAria II (BD). Whole lungs from E18, 19, 23, and 25 embryos and lung buds from E35 embryos were dissociated using *Bacillus licheniformis* protease (P5380, Millipore Sigma) with DNaseI. The live cells or EPCAM$^+$ cells were sorted using the following staining: rabbit anti-pig EPCAM antibody (MyBioSource, MBS2053510) and Donkey anti-rabbit IgG Alexa Fluor 647 (A-31573, Thermo Fisher Scientific); live or dead staining: SYTOX Green Nucleic Acid Stain (S7020, Thermo Fisher Scientific ), and Vybrant DycycleRuby Stain (V10309, Thermo Fisher Scientific). Cells were loaded on a 10x Genomics Chromium controller, aiming for 5,000 cell recoveries. For swine epithelium analyses, 4,464, 1,372, 5,528, and 5852 cells from E18, 19, 40, and 70 were recovered. For swine mesenchyme analyses, 12,288, 8,547, 11,006, 11, 851, 8, 478, and 7,596 cells from E18, 19, 23, 25, 35, 40, 70 were recovered.

#### Bulk RNA-seq analysis

For the cross-species bulk RNA-seq analyses, we input identifiers of a mouse (GSE122919; E12, 14, 16, 18, *n=3 each*), human (GSE121238; E10, 14, 20 weeks, *n=3 each*), and the swine lung development database (E26, 40, E70 distal, proximal, *n=3 each*). We annotated and mapped 12,424 common orthologues genes in mouse, human, and swine lung development using EggNOG and Homologene databases. To decide the swine lung stage, we generated a heatmap based on the normalized TPM value by comparing the human lung development database (GSE121238).

Briefly, we selected significant differentially-expressed genes (DEGs; Fold Change >1.5, FDR < 0.1) at each time point of human lung development using DESeq2 and applied the DEGs to the swine lung database. The normalization and unsupervised clustering of swine data were performed using Cluster3.0(*30*), and heatmaps were visualized using Java Treeview(*31*).

For the cross-species PCA, we used the TPM value of 12,424 common genes from mouse, human, and swine datasets in R ver 4.2.1(*32*). Among 12,424 common genes, PC1 minus-contributing genes were ranked by rotation of PC1, and the top 10% genes (1,242 genes) were further extracted to apply to gene ontology (GO) term enrichment analyses. GO enrichment analysis was used for each MCODE network to extract "biological meanings" from the network component. The top three best p-value terms were retained compared to all samples. We first identified all statistically enriched GO terms based on the default choices under Express Analysis using metascape. For generating the GO term network in Figure 2B, we selected a subset of representative terms from the full cluster and converted them into a network layout. Specifically, each term is represented by a circle node whose size is proportional to the number of input genes that fall under that term. Its color represents its cluster identity (i.e., nodes of the same color belong to the same cluster). Terms with a similarity score > 0.3 are linked by an edge (the thickness of the edge represents the similarity score). The network is visualized with Cytoscape with a "force-directed" layout and edge bundled for clarity. The same enrichment network has its nodes colored by p-value, as shown in the legend. Venn charts were drawn for 230 COSRP genes using the jvenn software(*33*).

**Cross-species scRNA-seq data analysis**

For the cross-species analyses, we used SCGEATOOL to combine datasets of the mouse (E9.5; GSE136689, E12.5; GSM4504959, E15.5; GSM4504960, E17.5; GSM4504961), human (E-MTAB-8221), and our swine lung development (E18, 19, 35, 40, 70 epithelium and E18, 19, 23, 26, 35, 40, 70 mesenchyme). Subsequently, UMAP and 3D tSNE analyses were unbiasedly calculated. We manually selected 45 genes highly expressed in humans and swine but not mice among the 230 COSRP genes in the 3D-tSNE plot. Among 45 genes, we identified 15 genes ubiquitously enriched either in swine and human lung mesenchyme or epithelium across the time, but not mice. For scGRN analyses, we applied 45 COSRP genes across the species at each developmental time point.

**scRNA-seq data analysis**

All scRNA-seq assays were performed with Illumina NovaSeq 6000. Reads alignment to Sscrofa11.1 pig reference genome and generation of gene expression matrix were processed using 10x genomics cellranger software version 6.1.2. For processing, integration, and downstream analysis, the Seurat package and SCGEATOOL were used. Cells were selected in the range of 2,000 to 9,000 mapped genes. Feature data were scaled using the Seurat ScaleData function. Non-linear dimension reduction was performed using uniform manifold projection (UMAP) using the Seurat RunUMAP process. The clustering was performed using the Louvain algorithm, and parameters were set empirically by detecting marker genes in each cluster. Pseudotime analysis was performed using Monocle 3 with the default parameter. The clusters associated with leukocyte, erythroid, endothelial, and neuronal cells were excluded from mesenchyme analysis. Differential expression for clusters was performed using the Seurat FindAllMarkers function using the

Wilcoxon rank test. Marker genes were calculated for each cluster against the cells within all the other clusters.

#### Genomic sequence alignment analysis

By the GO network analysis using cross-species bulk RNA-seq datasets, we found the condensed four GO nodes (ASSR, Regulation of neuron projection development, Regulation of nervous system development, and Axongenesis) with highly enriched networks. We defined those four GO term networks as COSRP, containing overlapping 230 genes. Within those 230 COSRP genes, 45 genes were enriched in human and pig lung development but not in mice, analyzed by cross-species 3D tSNE plot using SCSEGATOOL. Among 45 COSRP genes, 17 showed ubiquitous expression patterns with moderate or high levels in swine scRNA-seq Seurat analysis (**Data S2**). Since mouse *PRKG1* CDS and *human CCDC85B* TSS were not annotated in the Ensemble, we selected human CDS (coding sequence) or promoters (2,000 bp upstream of TSS) of 15 COSRP (*SEMA3C, NR2F1, B3GNT2, ENAH, EFNA5, ADNP, ADAMTS7, ANTXR1, B4GALT5, COL5A2, ID4, SRP54, TERF2IP, TET1,* and *ZFP36L2*) and 11 HK genes (*EIF4G2, HNRNPA2B1, HNRNPC, RBX1, SEC62, SON, YBX1, EIF3H, PFDN5, SNRPE,* and *YWHAE*) in the Ensemble. For comparing eight animals, *NR2F1* and *SNRPE* were excluded from the analysis because the homologous sequence was not annotated in shrew mouse and golden hamster. Human TSS information was identified from Eukaryotic Promotor Database. In the case of multiple promoters, the primary promoter for each gene was used for the following analysis(*34*). The homology (% identity: percentage of nucleotide sequence identity, and % Gaps) was calculated using a pair-wise comparison program with the Smith-Waterman algorism(*35*) by Snapgene (ver. 6.1.1) software(Gap open penalty: 10.0, Gap extend penalty: 1.0). For the interspecies promoter or CDS analysis, we compared the human CDS or promoter (2,000 bp upstream of TSS) sequence of COSRP genes or HK genes with the homologous region of each species, identified through comparative genomics alignment (text) or Nucleotide program in the Ensemble. The graphs were visualized, and statistical analyses were performed using Prism (ver. 9).

#### Immunostainings

Tissue samples were fixed in 4% paraformaldehyde and embedded in OCT compound, reacted with the primary antibodies (SOX2 (Invitrogen, 14-9811-82, 1:200), SOX9(R & D, AF3075, 1:200), NKX2-1(Abcam, ab76013, 1:100), Pro-SPC(Seven hills, WRAB-9337, 1:1000), Pro-SPB (Seven hills, WRAB-55522, 1:1000) overnight at 4 degrees in PBS, and washed in PBS, then incubated with the secondary antibodies conjugated with Alexa488 (Invitrogen, A21208, A21202, each 1:300), 568 (Invitrogen, A10042, A11057, each 1:300), or 647 (Invitrogen, A31573, 1:300), and Hoechst 33342 (Invitrogen, R37605, 1 drop in 500ul PBS) for 2 hours. Samples were mounted with PloLong Gold antifade reagent (Invitrogen, P37606). All images were captured with a 10x or 20x objective lens using confocal microscopy (Zeiss LSM 710).

#### Morphometric analysis for immunostainings

To determine the mean fluorescent intensity (MFI) of the SOX2 and SOX9 of the pig epithelial cells, all images were captured with the same settings captured by confocal microscopy (Leica LSM 710) (*n=3*). More than two thousand NKX2-1 positive cells from each sample were randomly selected, and each cell's nuclear MFI for SOX2-A488 and SOX9-A568 channels were calculated with the algorithm of Nuclei Count using Aivia software(Ver 10.5.1). The graphs were visualized using Prism (ver. 9).

**Study approval:** All experiments involving animals were performed according to the protocol approved by Columbia University Institutional Animal Care and Use Committee and USAMRMC Animal Care and Use Review Office (ACURO).

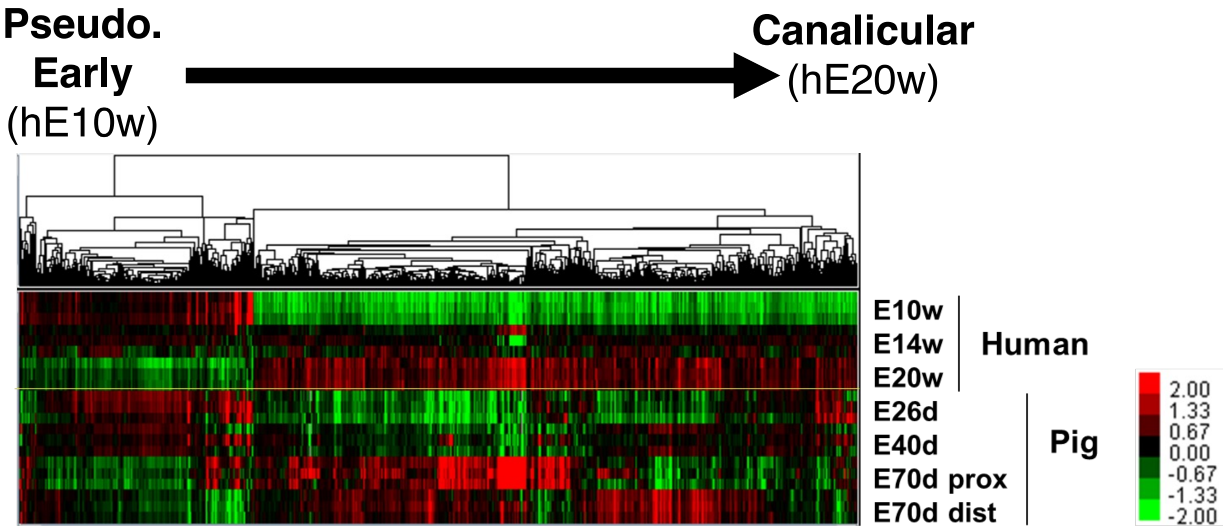


**Extended Data Figure 1. Swine lung developmental stage analysis by cross-species bulk RNA-seq** Heatmap of swine lung development at E26, E40 lungs, and E70 proximal or distal lungs depicting the top10% differentially expressed gene (DEG) found in human E10w, E14w, and E20w (gestation weeks 10, 14 and 20) during human lung development (data from GSE121238). The gene lists are in Supplementary Table 1. The hE10w-einrched genes were expressed at E26 and E40 swine lungs but low at E70, while the hE20w-enriched genes were enriched at both E70 proximal and distal regions.

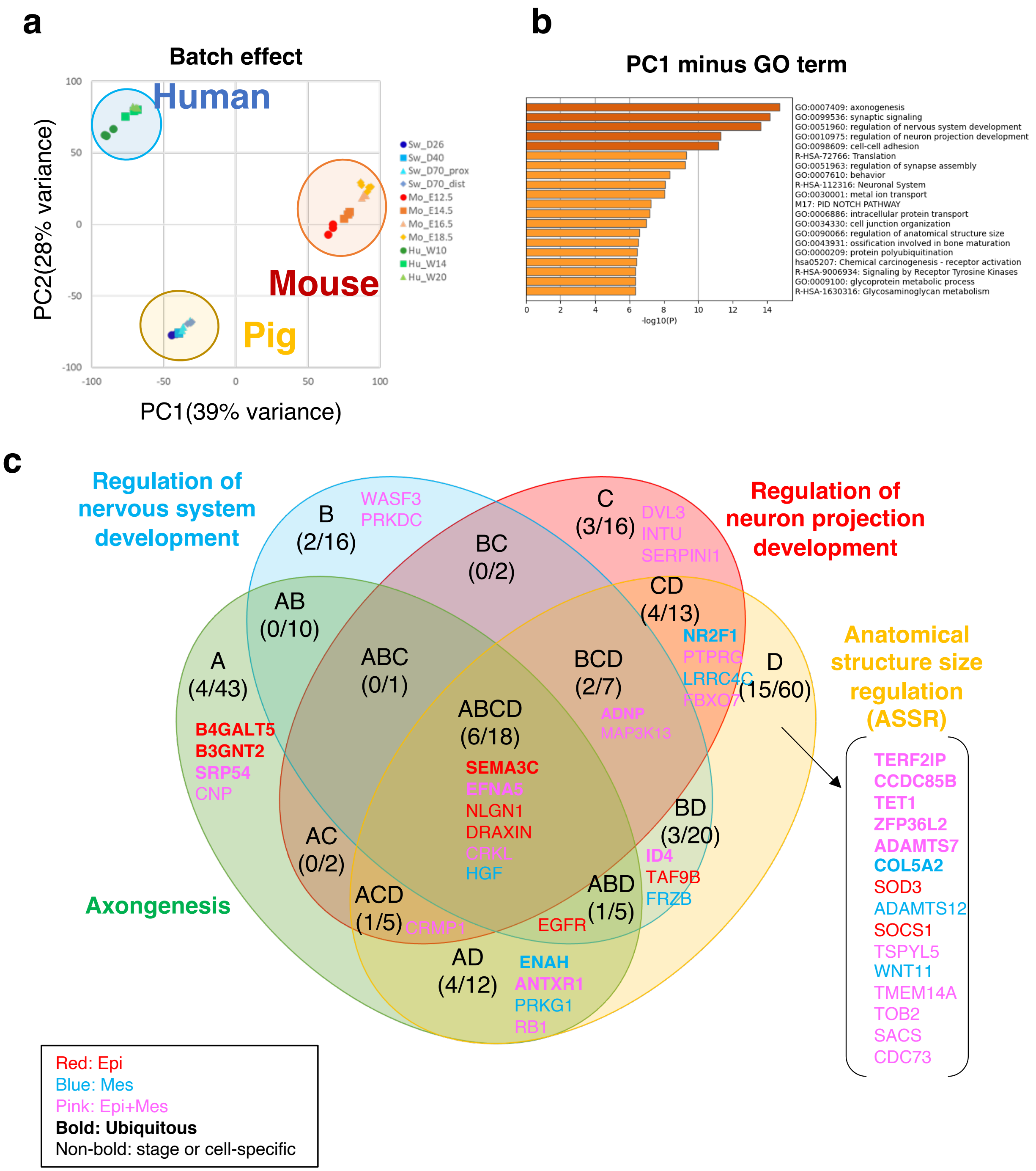


**Extended Data Figure 2. COSRP gene expression patterns during swine lung development (a)** PCA analysis of human, mouse, and pig embryonic lung samples showed batch effect. **(b)** Bar graph depicting GO terms contributing to PC1 minus axis (colored by p-values). **(c)** Venn diagrams: each COSRP gene classification to 4 GO terms (axonogenesis, regulation of nervous system development, regulation of neuron projection development, and ASSR). Gene names were colored by their expression pattern in pig scRNA-seq Seurat analysis. COSRP genes expressed in epithelial (red), mesenchymal (blue), or both epithelial and mesenchymal cells (pink). Bold font: ubiquitous expression throughout swine lung development. Non-bold font: stage-specific or cell type-specific expression pattern. The number in the Venn diagrams: the genes expressed in humans and pigs specifically but rarely in mice by the cross-species 3D-tSNE plot (left)/ the number of COSRP genes picked up by the cross-species bulk RNA-seq (right).

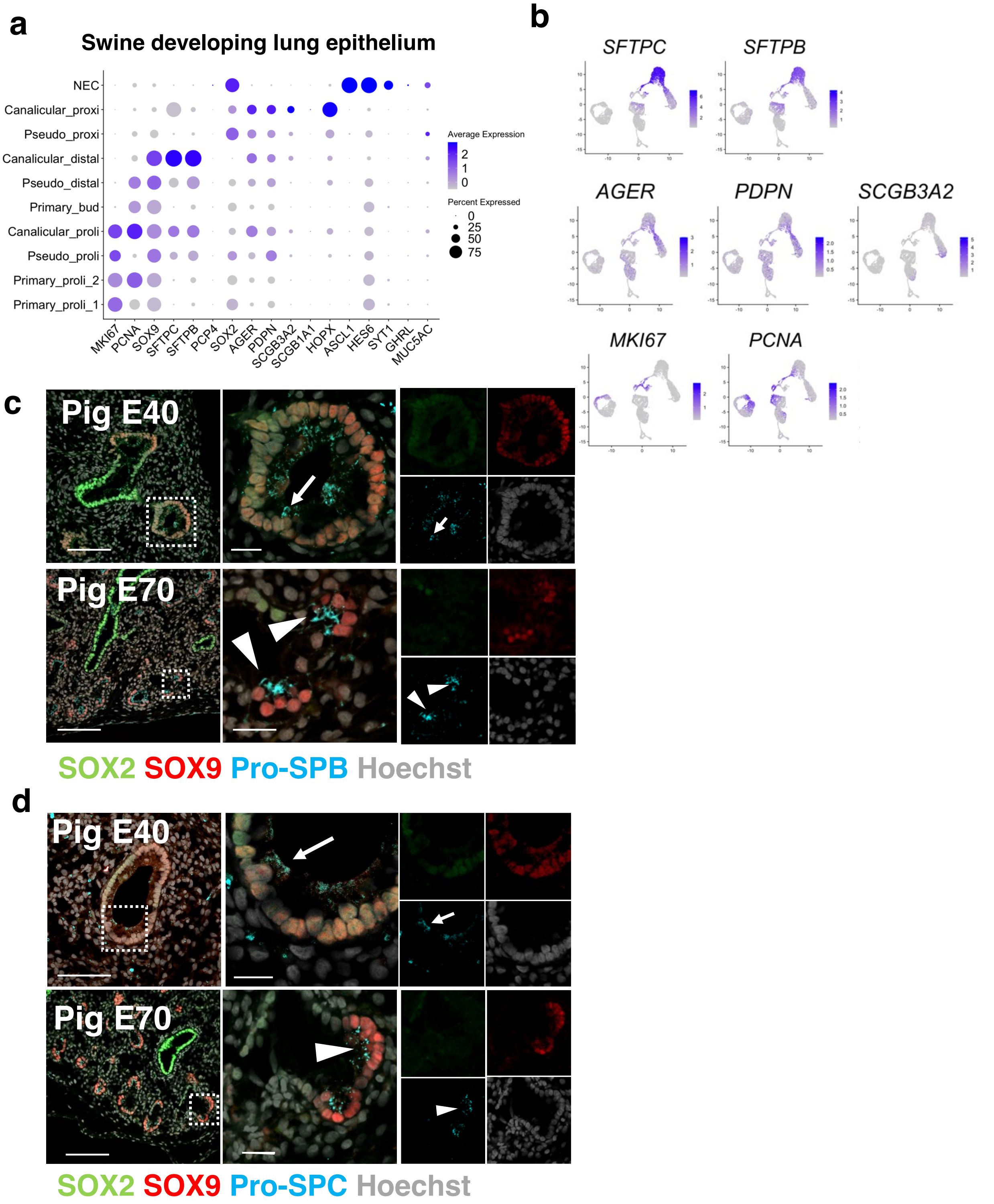


**Extended Data Figure 3. Expression patterns of mouse lung epithelial markers during pig lung development.** Dot plot (**a**) and UMAP feature plot (**b**) depicting the airway (*SCGB3A2*: secretory cells, *MUC5AC*: goblet cells), alveolar (*SFTPC*, *SFTPB*: type2 cells, *PDPN*, *HOPX*: type1 cells, bipotential progenitors), and proliferating cell markers (*MKI67*, *PCNA*) during pig lung development. Confocal IF imaging: SOX2 (green), SOX9 (red), and pro-SPB (cyan) (**c**) or pro-SPC (cyan) (**d**). Dense pro-SPB-positive or pro-SPC granules were present at E70 $SOX9^{+}SOX2^{-}$ sacculating cells (arrowheads) but less on the E40 $SOX9^{+}SOX2^{+}$ cells. Scale: left: 100µm, right: 20µm

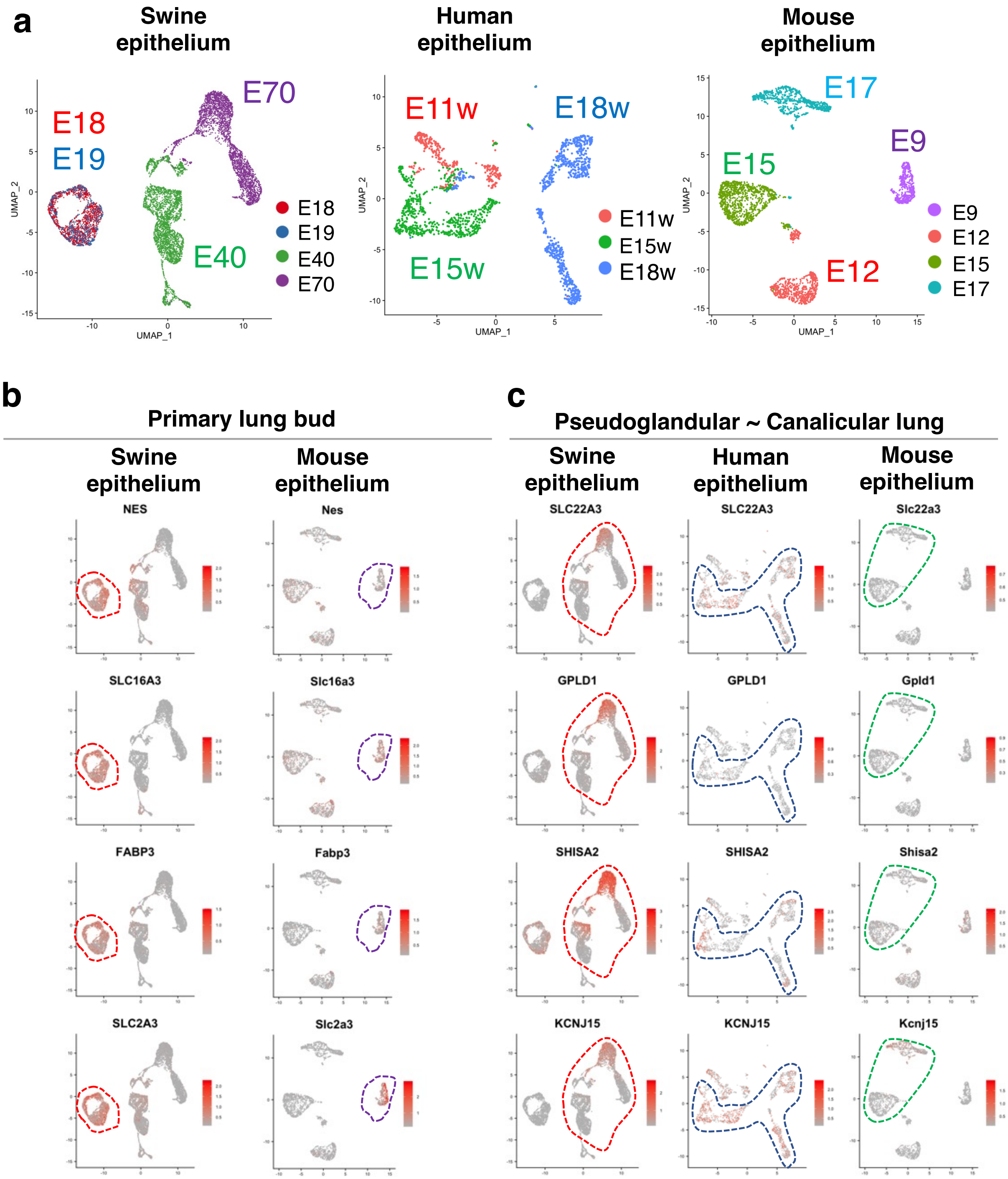


**Extended Data Figure 4. Identification of stage-specific lung bud markers during swine lung development (a)** UMAP visualization of each developmental time point in the swine, human, and mouse epithelial cells during lung development. **(b)** UMAP feature plot showing genes highly expressed at the pig primary lung bud stage (*NES*, *SLC16A3*, *FABP3*, and *SLC2A3*) compared to mouse lung development. **(c)** UMAP feature plot of genes highly expressed from pseudoglandular to the canalicular stage in pigs (*SLC22A3*, *GPLD1*, *SHISA2*, *KCNJ15*).

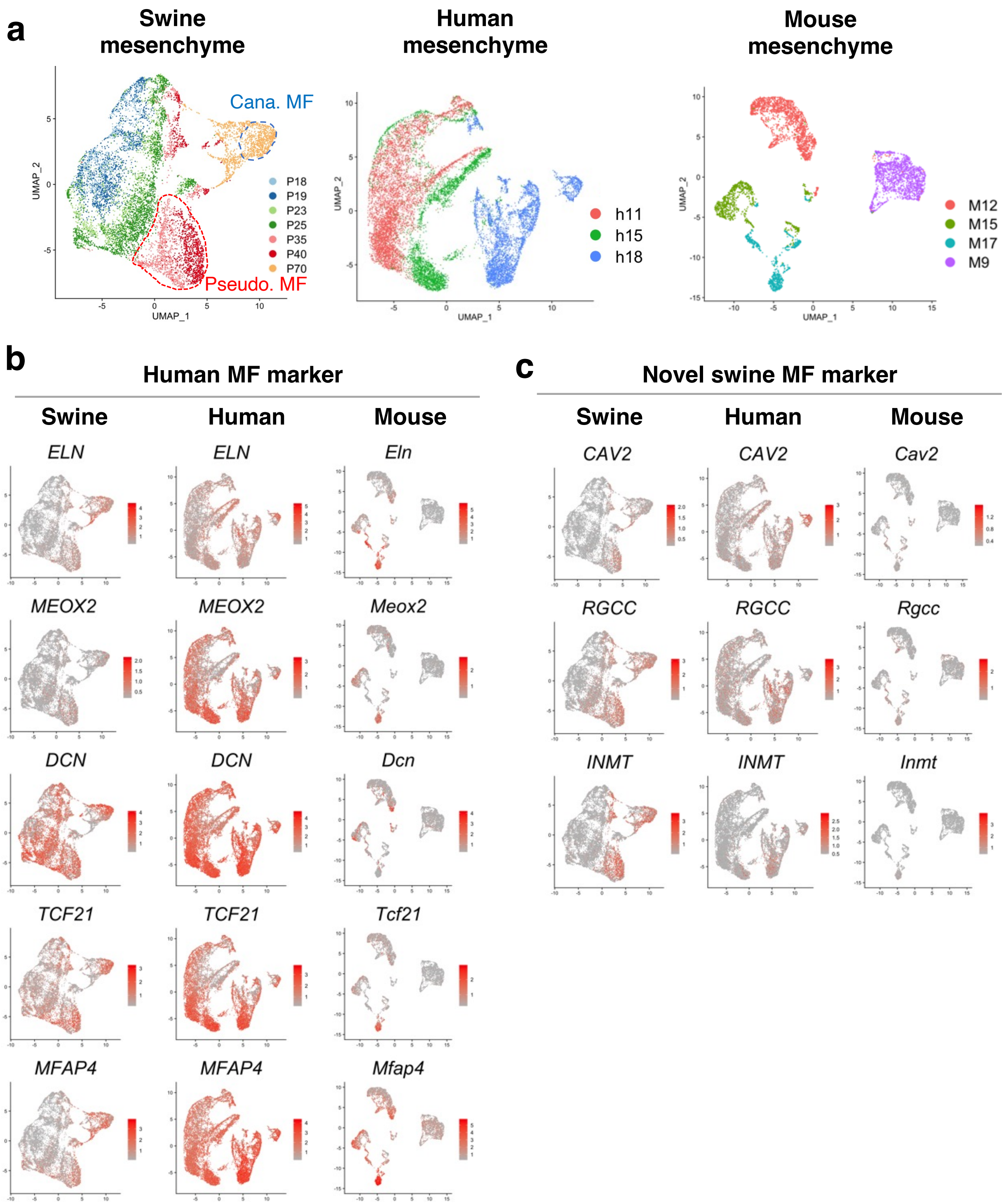


**Extended Data Figure 5. Expression patterns of matrix fibroblast (MF) associated genes in pig, human, and mouse lung development (a)** UMAP visualization of mesenchymal cells at each stage of lung development of pigs (pig gestation days: P18, P19, P23, P25, P35, P40, and P70), humans (human gestation weeks: h11, h15, and h18), and mice (mouse gestation days: M9, M12, M15, and M17). **(b)** Feature plots for human matrix fibroblast (MF) markers; *ELN, MEOX2, DCN, TCF21*, and *MFAP4*. During swine lung development, *ELN* and *MEOX2* enriched exclusively in MF. *ELN* was also expressed in SMC. During swine lung development, *DCN* and *TCF21* were not specific for MF but expressed in most lung mesenchyme over time except canalicular SMC. *MFAP4* was enriched in MF but also present in SMC, pericyte, and in a part of early pseudoglandular fibroblasts. (**c**) UMAP feature plots: novel MF genes enriched in swine and humans but not in mice: *CAV2, INMT,* and *RGCC* were enriched in MF, pericyte, proliferating pseudoglandular fibroblasts, and SMC. *RGCC* was also expressed in the early pseudoglandular fibroblast or chondrocyte.

**a**

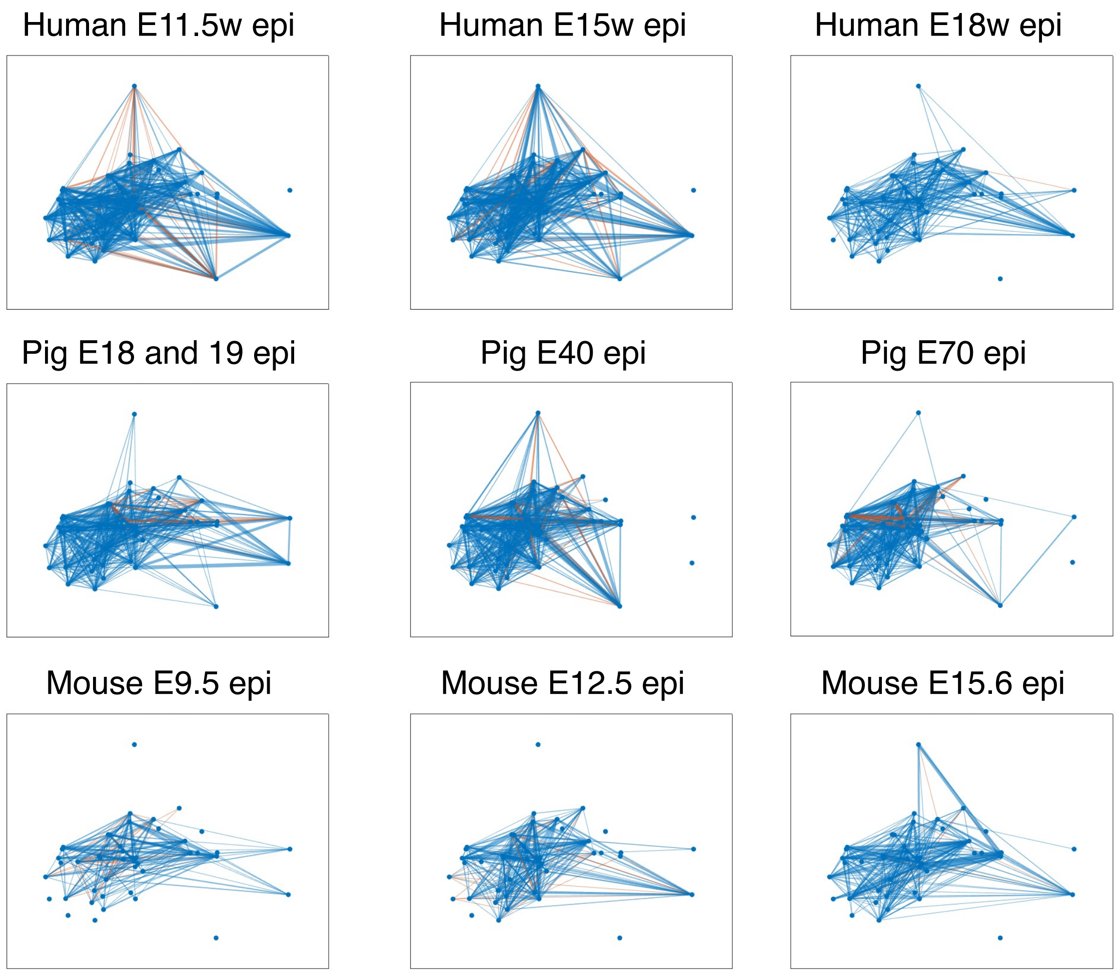
Human E11.5w epi
Human E15w epi
Human E18w epi
Pig E18 and 19 epi
Pig E40 epi
Pig E70 epi
Mouse E9.5 epi
Mouse E12.5 epi
Mouse E15.6 epi

**b**

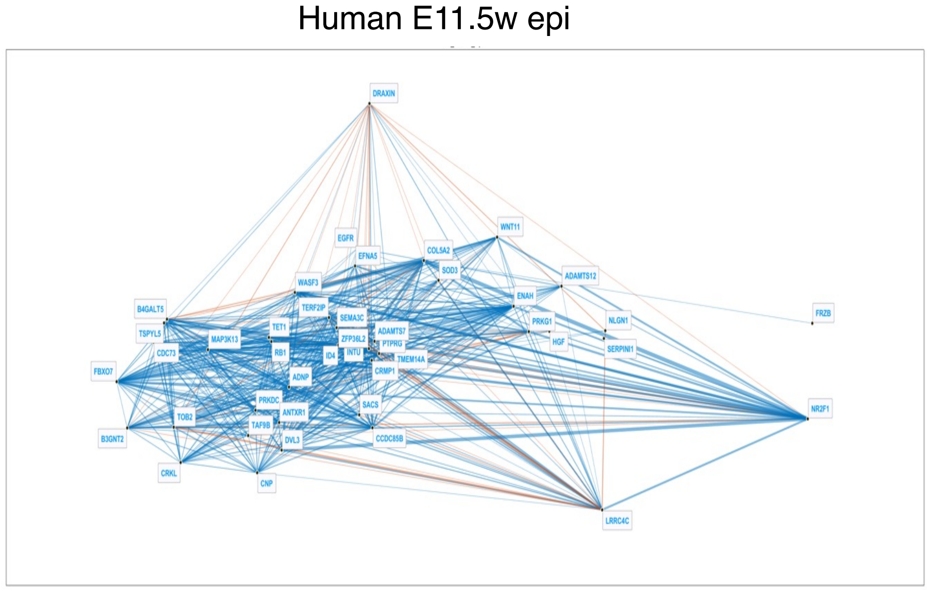
Human E11.5w epi
DRAXIN
WNT11
EGFR
EFNA5
COL5A2
SOD3
ADAMTS12
WASF3
ENAH
B4GALT5
TERF2IP
FRZB
SEMA3C
PRKG1
NLGN1
TSPYL5
TET1
ADAMTS7
MAP3K13
ZFP36L2
HGF
PTPRG
SERPINI1
CDC73
RB1
ID4
INTU
TMEM14A
FBXO7
CRMP1
ADNP
NR2F1
PRKDC
SACS
ANTXR1
TOB2
TAF9B
CCDC85B
B3GNT2
DVL3
CRKL
CNP
LRRC4C

**Extended Data Figure 6. Conserved COSRP Gene regulatory network (GRN) during human and swine lung development** (**a**) GRN visualization for COSRP genes in the epithelial cells at each stage of lung development of pigs (pig gestation days: P18, P19, P40, and P70), humans (human gestation weeks: h11, h15, and h18), and mice (mouse gestation days: M9, M12, M15). Over time, pig and human scGRN showed a highly dense network connected to the GRN nodule but sparse in mouse scGRN. (**b**) scGRN with gene annotation: *ZEP36L2, SEMA3C, TERF2IP, and ADNP* formed a central nodule in the scGRN of developing lung epithelium.

**Supplementary Table 1 (Separate file). List of DEGs in the human embryonic lung at 10w and 20w (GSE121238)**
**Supplementary Table 2 (Separate file). List of COSRP genes with their expression levels in developing swine lung epithelium and mesenchyme analyzed by Seurat feature map.**
